# PXPermute: Unveiling staining importance in multichannel fluorescence microscopy

**DOI:** 10.1101/2023.05.28.542646

**Authors:** Sayedali Shetab Boushehri, Aleksandra Kornivetc, Domink J. E. Waibel, Salome Kazeminia, Fabian Schmich, Carsten Marr

**Affiliations:** Institute of AI for Health, Helmholtz Zentrum München - German Research Center for Environmental Health, Neuherberg, Germany; Institute of Computational Biology, Helmholtz Zentrum MünchenMunich - Helmholtz Munich - German Research Center for Environmental Health, Neuherberg, Germany; Technical University of Munich, School of Life Sciences, Weihenstephan, Germany; University of Hamburg, Department of Informatics, Hamburg, Germany; Technical University of Munich, Department of Mathematics, Munich, Germany; Data & Analytics, Pharmaceutical Research and Early Development, Roche Innovation Center Munich (RICM), Penzberg, Germany

**Keywords:** channel importance, staining importance, interpretable artificial intelligence, Image flow cytometry, deep learning, machine learning, computer vision, cell profiling

## Abstract

Imaging Flow Cytometry (IFC) enables rapid acquisition of thousands of single-cell images per second, capturing information from multiple fluorescent channels. However, the conventional process of staining cells with fluorescently labeled conjugated antibodies for IFC analysis is labor-intensive, costly, and potentially detrimental to cell viability. To streamline experimental workflows and reduce expenses, it is imperative to identify the most relevant channels for downstream analysis. In this study, we present PXPermute, a user-friendly and powerful method that assesses the significance of IFC channels for a given task, such as cell profiling. Our approach evaluates channel importance by permuting pixel values within each channel and analyzing the resulting impact on the performance of machine learning or deep learning models. Through rigorous evaluation on three multi-channel IFC image datasets, we demonstrate the superiority of PXPermute in accurately identifying the most informative channels, aligning with established biological knowledge. To facilitate systematic investigations of channel importance and aid biologists in optimizing their experimental designs, we have released PXPermute as an easy-to-use open-source Python package.

## Introduction

Imaging flow cytometry (IFC) is a high-throughput microscopic imaging technique that captures multiparametric fluorescent and morphological information from thousands of single cells. This versatile method allows researchers to rapidly record and analyze large cohorts of cells, providing valuable insights into cell populations ^1–3^. IFC has been used for profiling complex cell phenotypes and identifying rare cells and transition states ^3^, making it an indispensable tool for various applications such as drug discovery ^4^, disease detection, diagnosis ^3^, and cell profiling ^5–8^. Fluorescent staining of cells, while informative, is not without its limitations. The panel design process for imaging flow cytometry (IFC) can be time-consuming and expensive ^9^. Moreover, the use of multiple stains can introduce complications such as spectral overlaps and compensation issues ^9^. Additionally, fluorescent staining has the potential to harm cells, and artifacts may arise during the staining and sample preparation steps ^4^. To address these challenges, it is crucial to carefully select a restricted number of stainings, simplifying laboratory procedures, reducing costs, preserving cell integrity, and enabling the evaluation of new fluorescent stainings. This is particularly significant in the context of delicate cells in hematological diseases ^3^ or synapse formation ^10^. Machine learning methods, such as image classification, promise to deliver accurate, consistent, fast, and reliable predictions ^11–13^. A few open-source libraries have recently been published, specializing in machine learning for IFC analysis ^10,11,14,15^. These libraries provide in-model interpretability like random forest ^16^ or post-model interpretability using methods such as Grad-CAM ^17^ for convolutional neural networks. However, none of them are designed specifically to evaluate the importance of fluorescent channels and require adaptation to address this specific task.

A natural solution to identify the most important fluorescent stainings is to systematically remove single or combinations of channels and retrain and re-evaluate the model to investigate the channel’s importance for the model’s performance. However, such an approach is time-consuming and computationally costly, requiring 2^*n*^ models to be trained, where *n* is the number of channels. Another solution is to design model architectures that allow for the analysis of different channel features separately. In that direction, Kranich et al. ^18^ proposed a convolutional autoencoder (CAE) in which each channel is embedded separately using an encoder and one shared decoder. The model embeddings are then concatenated and passed to a random forest classifier that enables channel importance rankings, as each channel is encoded separately. This method has two main limitations: (i) It is bound to the specific model architecture, and training the CAE is only manageable if the number of channels is limited, as training many separate encoders can be computationally very expensive; (ii) Feature correlations across channels are ignored by extracting features from each channel separately, which can negatively affect model performance or miss meaningful correlations among stainings ^19,20^.

We have developed PXPermute, a model-agnostic post-model interpretation method that identifies channel importance, ranks channels according to their contribution to a downstream task’s performance, and improves upon the previously mentioned shortcomings. We compare PXPermute with adapted state-of-the-art post-model interpretability methods on three publicly available IFC datasets. Our work identifies the most important channels that align with each dataset’s biology. PXPermute can also identify the least informative stainings, which might be eliminated from the experiment without affecting model performance. To the best of our knowledge, PXPermute is the first method that systematically studies channel importance and can lead to optimized workflows in multichannel fluorescent staining experiments.

## Results

### PXPermute - A model-agnostic post-hoc interpretability method for channel importance

We propose PXPermute, a model-agnostic post-hoc interpretability method for channel importance (Fig. 1a). Let’s assume there exist a dataset with images *x* with *n* channels. Moreover, for each *x* there is a corresponding label *y* covering *m* classes. Finally, a deep learning or classical machine learning model *f*(*x, y*) is already trained on images *x* and label *y*. To apply PXPermute, for each class (*cl*) in the dataset, the metric *M*_*cl*_ (e.g. accuracy) is evaluated on an independent test set. Next, for each channel *Ch*, PXPermute randomly shuffles pixels, and the model *f*(*x*^*Ch*^) calculates the new performance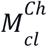 after the disturbance to the input image *x*^*Ch*^. This process is done iteratively for each channel *Ch*. We define the difference between the original performance with the new performance as: 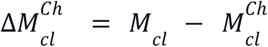 The channel importance for a channel *Ch*can be obtained by aggregating (e.g., average) the differences for each class 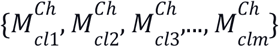 (Fig 1b, and Methods).

**Figure 1:**
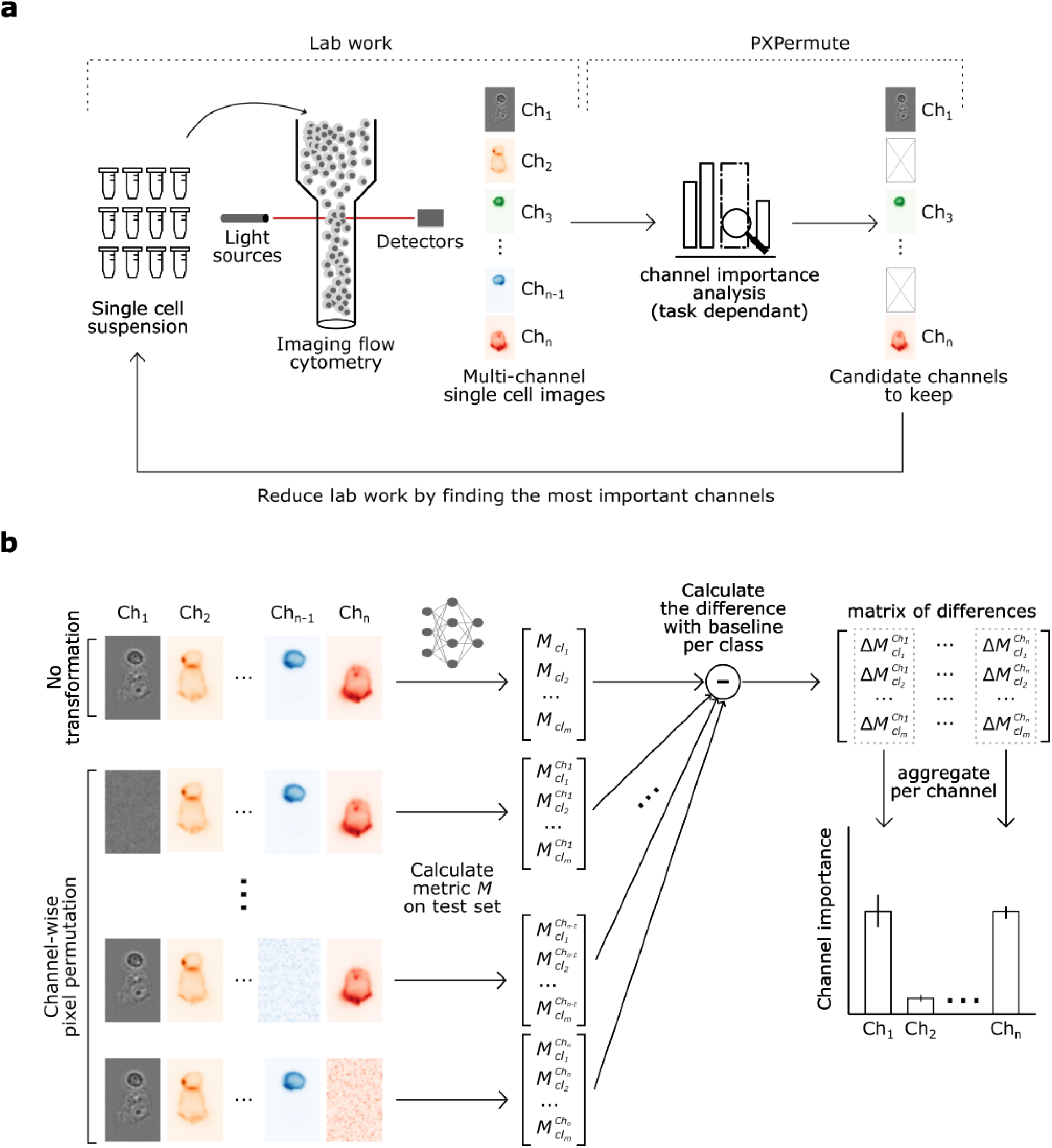
PXPermute allows to identify most important fluorescent channels in a multi-channel imaging flow cytometry experiment and thus reduces lab work and expenses. **a.** Schematic of a PXPermute embedded end-to-end analysis. In the first part, biologists image thousands of single cells using imaging flow cytometry in different fluorescent and brightfield channels. The second part PXPermute, will select the most important channels based on the task. **b**. Schematic of PXPermute, a simple yet powerful method to find the most important channels. In the first step, a performance metric *M*_*cl*_ (such as accuracy or F1-score) is calculated per class *cl*. Then each channel *Ch* is shuffled, and the performance per class *cl* is calculated as 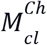 Finally, the difference between the original performance and the permuted one is calculated. These differences are averaged and yield the channel importance.

We selected three publicly available IFC datasets to demonstrate PXPermute’s potential to rank fluorescent stainings as well as stain-free channels in multichannel images. The first dataset contains 15,311 images of two classes, comprising 8,884 apoptotic cells and 6,427 non-apoptotic cells with only a bright field and one fluorescent channel (Fig. 2a and Methods). The second dataset contains 5,221 images of lymphocytes that form immunological synapses ^10,21^. Images contain a single cell, two cells, or more than two cells. It is distributed into the following nine classes: B cells, T cells without signaling, T cells with signaling, T cell with smaller B cells, B & T cells in one layer, synapse without signaling, synapse with signaling, no cell-cell interaction, and multiplets. This data set includes eight channels: brightfield, antibody (Ab), CD18, F-actin, MHCII, CD3, P-CD3ζ, and Live/Dead stainings (Fig. 2b and Methods). Considering that the dataset consists of live T or B cells with no antibodies, the Ab and Live/Dead channels do not contain any relevant information. Therefore they are used as a sanity check in the channel importance analysis. Finally, the white blood cell dataset ^12^ includes 98,013 images with eight classes and twelve channels. The classes are CD14+ monocyte, CD15+ neutrophil, CD19+ B cell, CD4+ T cell, CD56+ NK, CD8+ T cell, NKT, and Eosinophil. The channels include brighfield1 (BF1), CD15, SigL8, CD14, CD19, darkfield (DF), CD3, CD45, brightfield 2 (BF2), CD4, CD56, and CD8 (Fig. 2c and Methods).

**Figure 2:**
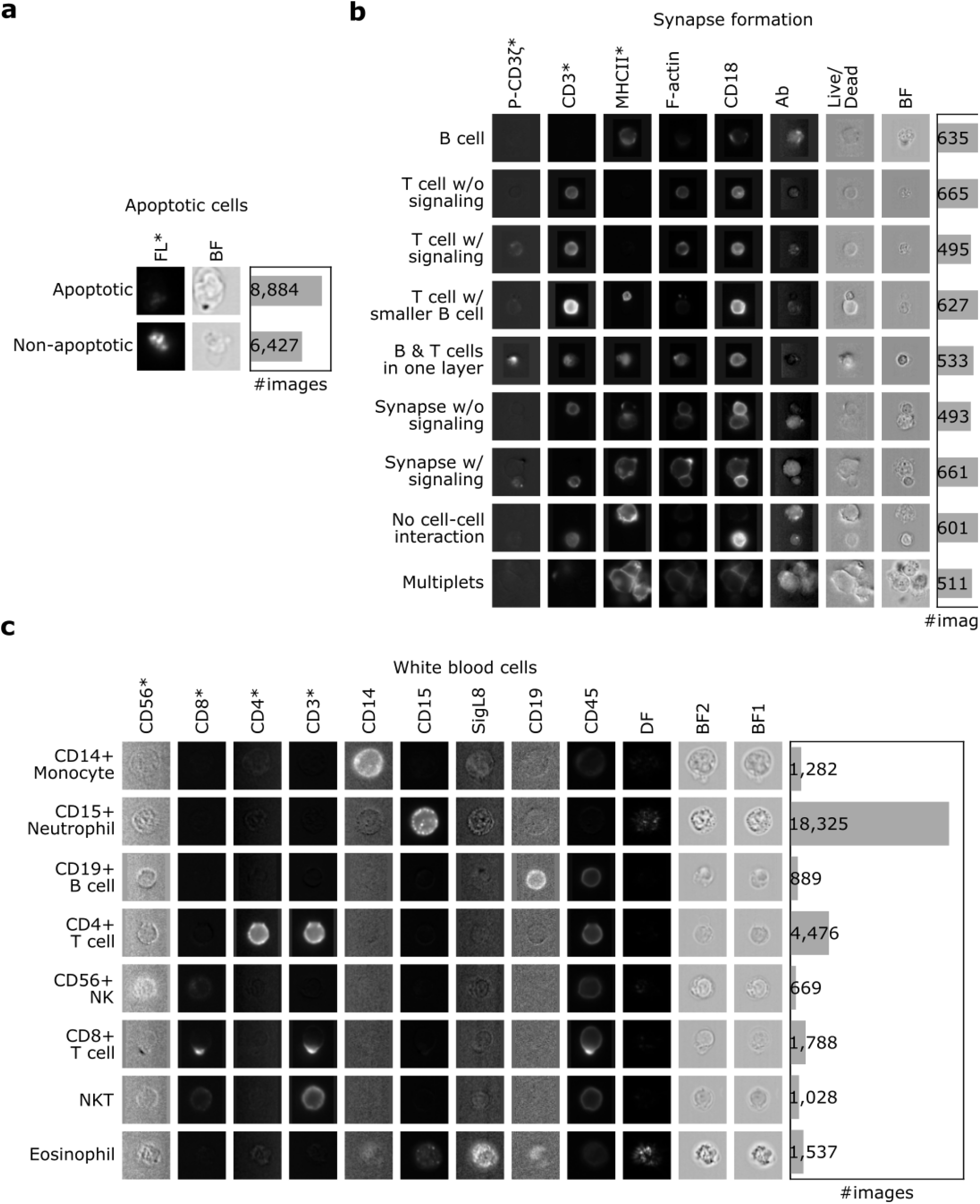
Three datasets with various numbers of images and channels are used to evaluate PXPermute. Rows indicate classes, and columns indicate the channels. Channels marked with (*) indicate that those channels have been identified as the most important channels in previous works. **a** Apoptotic cells: a dataset containing 15,311 images with one stain-free, bright-field (BF) and a fluorescent channel (FL). **b** Synapse formation: a dataset containing 5,221 images with one stain-free channel (BF) and seven fluorescent channels, namely Antibody (Ab), CD18, F-actin, MHCII, CD3, P-CD3ζ and Live/Dead. **c** White blood cells: a dataset containing 2,9994 images with three stain-free channels, two bright fields (BF1 & BF2), a dark-field (DF), and nine stained channels including CD15, SigL8, CD14, CD19, CD3, CD45, CD4, CD56, and CD8.

### PXPermute detects the most important channels in alignment with biological knowledge

As a proof of concept, we used PXPermute on a ResNet18 ^22^ pretrained on ImageNet ^23^, a widely used deep learning model for image classification ^24,25^. Considering that this model was designed originally for natural images with three channels, we replaced the first layer of the network by matching the number of channels per dataset. Also, the classification layer was changed based on the number of classes in each dataset (see Methods for details). We trained the ResNet18 on each dataset to classify the cells into the given cell types and compared it to the state-of-the-art performance. For the apoptotic vs. non-apoptotic cells, the model reached the performance of F1-macro = 0.97±0.01 (mean and standard deviation from five-fold cross-validation). For the synapse formation data set, the model reached an F1-macro performance of 0.95±0.01. Finally, for the white blood cell dataset, our model reached 0.98±0.01 of F1-macro (Methods).

Considering that no method has been previously developed for assessing channel importance, we adapted already existing methods used for pixel-wise interpretation, including occlusion ^26^, DeepLift ^27^, Guided Grad-CAM ^17^, integrated gradients ^28^, and LRP ^29^ (Table 1 and Methods). We modified the occlusion method to occlude the images channel-wise so that the whole channel is replaced with the 0. Similar to PXPermute, the drop in model performance is considered as the channel importance (Supplementary Fig. 1a). The other methods originally provide pixel-wise importance for each image, typically visualized as a heatmap ^30^. We aggregate the pixel importance per channel to adapt them to calculate a channel-wise importance score (Supplementary Fig. 1b). Except for channel-wise occlusion, all these methods are not model agnostic and strongly depend on the quality of the pixel importance estimation and its aggregation to obtain a channel score.

**Table 1:** Adaptation of used methods for benchmarking. Apart from PXPermute, designed for channel importance, all other methods are adapted from their original design. Their other advantages and disadvantages are based on model specificity, design complexity, run-time, and the need for tuning.

| Method name | Adaptation | Advantages | Disadvantages |
| --- | --- | --- | --- |
| PXPermute | None | + Open source<br>+ Model agnostic<br>+ Simple concept<br>+ Keeps the data distribution<br>+ No parameter tuning | - Higher run time |
| Channel-wise occlusion | Occluding channels instead of pixels | + Rapid run-time<br>+ Model agnostic<br>+ Simple concept<br>+ No parameter tuning | - Changes the data distribution |
| Guided GradCAM (Selvaraju et al.) | Using the median of the values per channel | + Rapid run-time<br>+ Open source | - Model-specific<br>- Requires parameter tuning<br>- Needs adaptation |
| Integrated gradients (Sundararajan et al.) | Using the median of the values per channel | + Rapid run-time<br>+ Open source | - Model-specific<br>- Requires parameter tuning<br>- Needs adaptation |
| DeepLift (Shrikumar et al.) | Using the median of the values per channel | + Rapid run-time<br>+ Open source | - Model-specific<br>- Requires parameter tuning<br>- Needs adaptation |
| LRP (Bach et al.) | Using the median of the values per channel | + Rapid run-time<br>+ Open source | - Model-specific<br>- Requires parameter tuning<br>- Needs adaptation |

We benchmarked all interpretation methods on the trained models in the next step. With each model training in the cross-validation scheme, we also run each interpretation method. Therefore we have five values per method for each channel. To provide a comparative overview, we normalized each channel’s importance to the interval of 0 and 1 (Fig. 3). Hence, the most important channel obtained an importance score of 1, and the least important channel had a score of 0. For simplicity, the number of existing channels in the datasets as well as previous relevant works, we only focus on the top-1 channel for the apoptotic cell dataset and the top-3 channels for the synapse formation and the white blood cell datasets.

**Figure 3:**
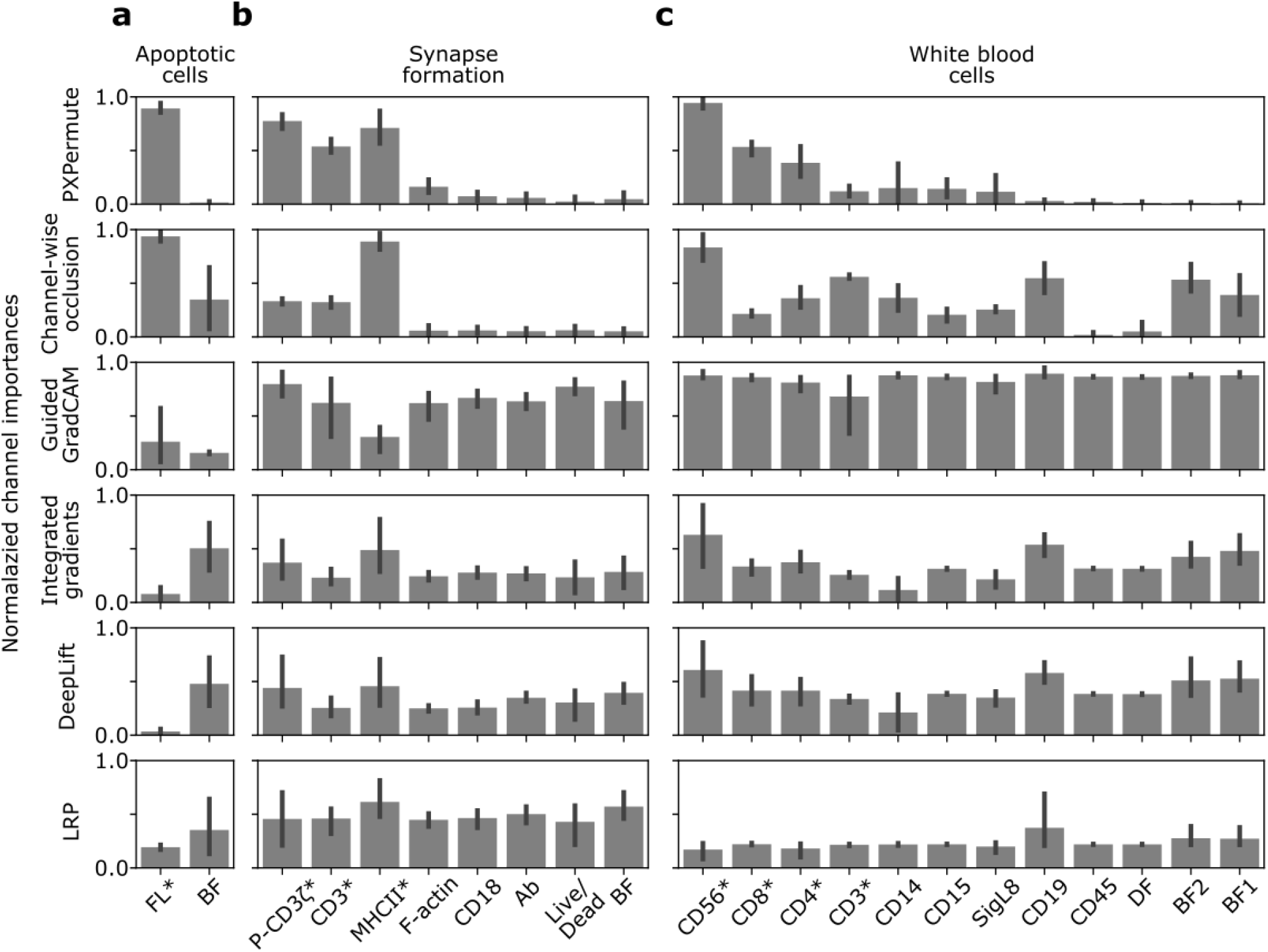
PXPermute robustly identifies the most important channels. Each dataset’s channel importance is depicted and normalized between zero and one. The higher the values, the more important the channels. The error bars are based on a five-fold cross-validation scheme. Channels marked with (*) indicate that those channels have been identified as the most important channels in previous works.

We first applied the channel importance methods to the apoptotic cells dataset. Kranich et al. previously showed on this dataset that the fluorescent channel is more important than the brightfield channel for predicting apoptotic cells ^18^. PXPermute, channel-wise occlusion, and Guided GradCAM correctly identified the fluorescent channel as the most important channel for predicting apoptotic cells, which aligns with previous work ^18^. In contrast, integrated gradients, DeepLift, and LRP identify brightfield as the most important channel (Fig. 3a and Supplementary Fig. 2a), leading to a false conclusion on this dataset.

For the synapse formation dataset, we previously showed that the most important channels are CD3, MHCII, and P-CD3ζ ^10,21^. PXPermute identifies the top channels P-CD3ζ, CD3, and MHCII in line with previous knowledge (Fig. 3b and Supplementary Fig. 2b). Channel-wise occlusion finds the same top-3 channels but in a different order. GradCAM identifies P-CD3ζ, Live/Dead, and CD18 as the most important channels. This order does not fit to our prior knowledge because Live/Dead is an irrelevant channel, as the original authors only annotated the live cells. Integrated gradients and DeepLift identify CD3, P-CD3ζ, and BF as the most important channels. Finally, LRP identifies CD3, BF, and Ab as the most important channels. This contradicts our knowledge, as no antibody exists in the dataset.

The white blood cell dataset has the largest number of channels and the highest variation of the channel rankings. This dataset had not been used for channel importance before. However, from the work of Lippeveld et al. ^12^, it is possible to infer that CD3, CD4, CD56, and CD8 are the most important channels. PXPermute suggests that the top-3 channels are CD56, CD8, and CD4, confirming Lippeveld et al.’s work. Channel-wise occlusion suggests that CD56, CD3, and CD19 are the most important channels. Guided GradCAM suggests that BF1, CD56, and BF2 are the most important channels. Integrated gradients identified CD46, CD19, and BF1 as the most important channels. DeepLift indicates that the most important channels are CD56, CD19, and BF1. Finally, LRP implies that CD19, BF2, and BF1 are the most important channels (Fig. 3c and Supplementary Fig. 2c). Our method identified channel importance closely aligned with existing baselines and expert findings ^10,12,18^.

### Identifying and removing unnecessary channels with PXPermute

To validate channel importance rankings suggested by PXPermute and the other methods, we applied the remove-and-retrain principle ^31^: First, we sorted image channels according to their predicted importance score in ascending manner, from the least important channel to the most important. Then we iteratively removed channels from the dataset, from the least important to the most important channel. Secondly, after each removal, the classification model was retrained on the dataset containing the subset of channels to perform the same classification task as before. During the remove-and-retrain process, we fixed model hyperparameters. We repeated this procedure for every channel ranking (see Fig. 3). In an ideal scenario, removing a channel that is not important for the model prediction should not affect model performance while removing a channel that is important for the model should highly affect its performance. Therefore, if the performance of a model during the remove-and-retrain process drops faster for a given sequence of channels than for another sequence of channels, it shows that the sequence ordering was not optimal. To numerically compare the drops in classification performance, we calculate the average F1-macro across all the runs (see Fig. 3).

For the apoptotic cells dataset, the order suggested by PXPermute, channel-wise occlusion, and Guided GradCAM was the optimal sequence. They showed the highest performance (average F1-macro of 0.97±0.01, n=10 based on five-fold cross-validation and two channels). They were followed by LRP (0.83±0.14), integrated gradients (0.83±0.15), and DeepLift (0.82±0.15) (Fig. 4a).

**Figure 4:**
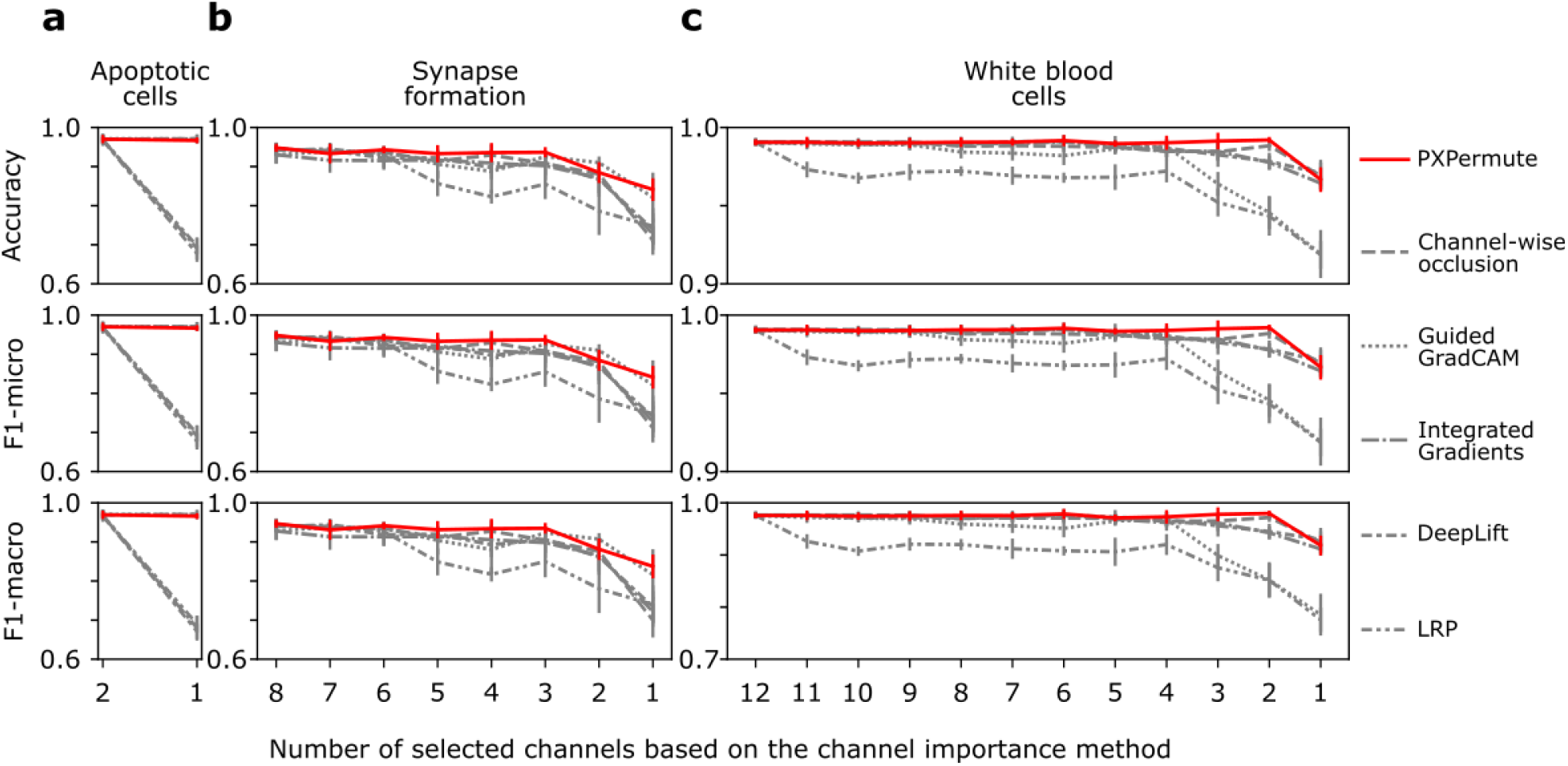
PXPermute outperforms other methods in identifying the correct channel ranking based on a remove-and-retrain procedure. The remove-and-retrain based on the channel ranking is performed on the apoptotic cells (2 channels), synapse formation (8 channels), and white blood cells (12 channels) datasets. In each case, the channels are ascendingly sorted according to their predicted importance score, from the least important channel to the most important. Then the channels are iteratively removed from the dataset, from the least important to the most important channel. After each removal, the classification model was retrained on the dataset containing the subset of channels to perform the same classification task as before. Methods with better rankings would stay higher throughout the plot. PXPermute performed better in finding the optimal channel rankings than other methods.

For the synapse formation dataset, PXPermute predicted the best sequence of channels (average F1-macro=0.92±0.04, n=40 based on five-fold cross-validation and eight channels). It was followed by GuidedGradCAM (0.90±0.05), DeepLift (0.89±0.07), channel-wise occlusion (0.89±0.08), integrated gradients (0.88±0.07), and LRP (0.86±0.08) (Fig. 4b).

For the white blood cell dataset, PXPermute, channel-wise occlusion, and DeepLift predicted the best sequence of channels (average F1-macro=0.97±0.02, n=60 based on five-fold cross-validation and 12 channels). It was followed by integrated gradients (0.96±0.02), Guided GradCAM (0.94±0.06), and LRP (0.90±0.05) (Fig. 4c). The same pattern could be observed for average F1-micro and accuracy. Therefore, PXPermute is the only method that correctly detects the order of the importance of the channels in all datasets (Fig. 4).

### PXPermute finds the optimal panel with a minimal number of stainings

Identifying a minimal set of stainings required for a downstream task has manifold advantages, such as the potential to reduce staining artifacts, save cost and time in sample preparation. It also allows adding new, more meaningful stainings to the panel. Therefore, identifying the minimum set of channels a model requires to deliver optimal performance is highly beneficial. To investigate this effect, we compared the model’s performance trained with only non-stained channels, with only the most important channels according to PXPermute adding non-stain channels and with all channels. We conducted these experiments on the synapse formation and the white blood cell dataset (Fig. 5) and skipped the apoptotic cell dataset, as it has only two channels.

**Figure 5:**
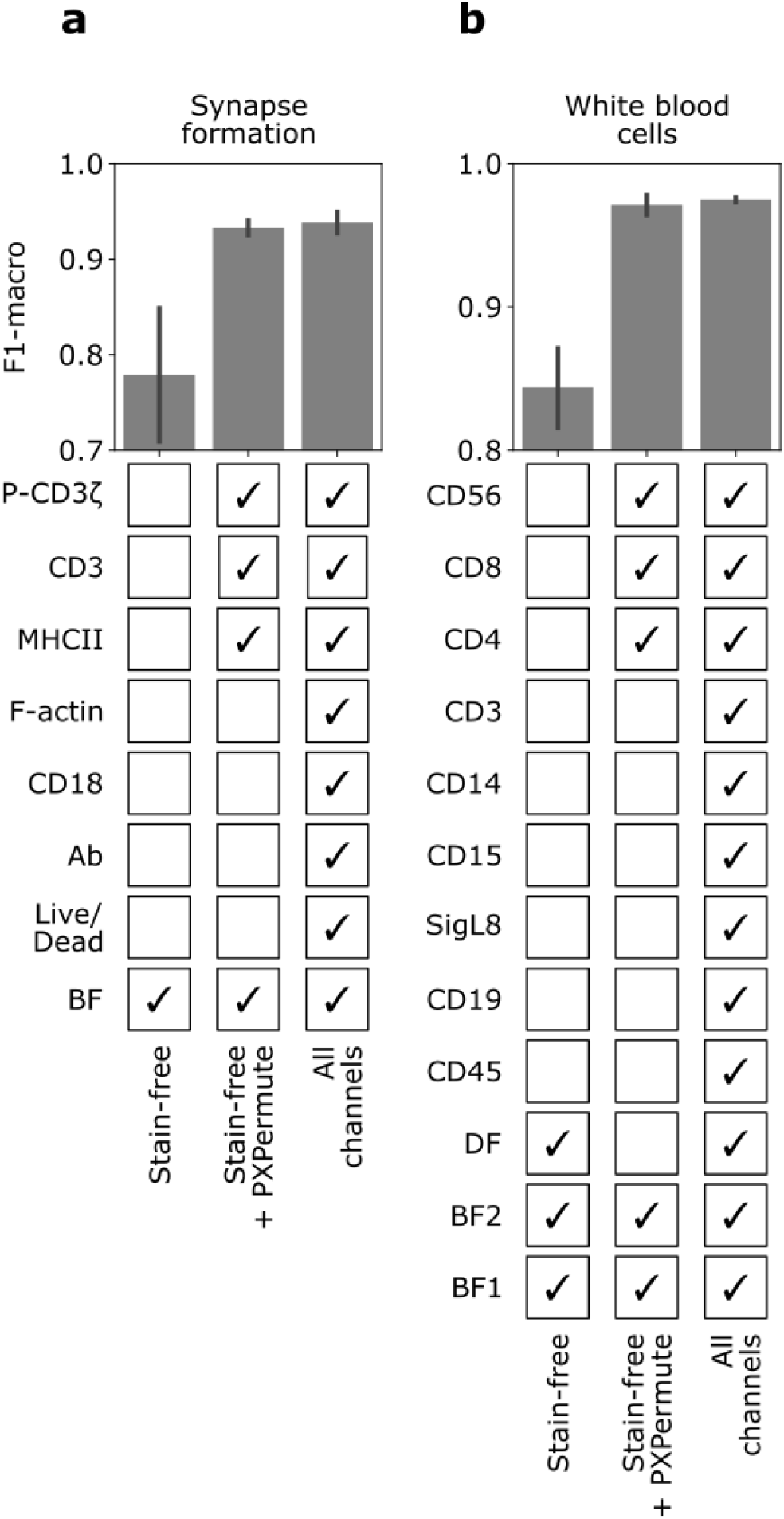
PXPermute finds an optimal channel selection that performs similarly to using all the channels. For the synapse formation (a) and white blood cells datasets (b), the model was trained on the stain-free channels (lower bound), stain-free + top-3 channels identified by PXPermute, and all channels (upper bound). PXPermute rankings lead to fewer stainings (3 out of 7 in a, 3 out of 8 in b) without a significant loss in performance.

For the synapse formation, the stain-free selection of channels (BF) achieves 0.78±0.09 F1-macro in a five-fold cross-validation setup. Adding the suggested channels from PXPermute, namely, MHCII, CD3, and P-CD3ζ, improves the classification to 0.93±0.01 F1-macro, which is not so different from 0.94±0.01 F1-macro, which is the accuracy when using all channels. For the white blood cell dataset, the stain-free selection of channels (BF1, BF2, and DF) achieves 0.84±0.04 F1-macro. By adding only the three most important channels identified by PXPermute, namely CD4, CD56, and CD8, the classifier archives the performance of 0.97±0.01 F1-macro, which is not significantly less than a model trained on all channels, reaching 0.97±0.00 F1-macro. We have shown that PXPermute can identify the most important channels for a cell classification task. The required channels can be halved without significantly decreasing the model’s performance.

## Discussion

In this study, we introduced PXPermute, a pioneering post-model interpretability method that assesses the impact of fluorescent stainings on cell classification. By applying PXPermute to three publicly available imaging flow cytometry (IFC) datasets, we demonstrated its efficacy in accurately identifying the most significant channels in accordance with biological principles. Moreover, PXPermute facilitated panel optimization by recommending a reduced number of stainings that maintained comparable performance to using all channels.

Channel importance in the context of staining has received limited attention in previous studies, with only a handful of works touching upon its significance ^10,18^. Therefore, our research represents the first comprehensive investigation into systematically exploring channel and staining importance. Kranich et al. proposed a convolutional autoencoder that can rank the channels after training a random forest on the decoupled features of each encoder ^18^. However, this work is limited as it is constrained to a specific model architecture and dataset with the smallest number of channels. Previously, we performed a remove-and-retrain ablation study to identify the most important channels ^10^. However, this method needs a clearer channel ranking, and the computational expense of retraining the model with all possible channel combinations is prohibitive. PXPermute addresses these limitations and provides a model-agnostic method for feature ranking that can be applied independently of the number of channels or model architecture.

To prove PXPermutes’ robustness, we studied it on three different publicly available datasets. We identified that the fluorescent channel is more important than the brightfield channel for the apoptotic vs. non-apoptotic cells, which aligns with the previous work ^18^. In the synapse formation study, MHCII, CD3, and P-CD3ζ were identified to detect immunological synapses, again in alignment with the original study ^10^. While other methods also found a high-performing combination of channels, none aligned with underlying biology, and they were probably only hinting at possible artifacts in the dataset. Finally, for the white blood cell dataset, the authors did not provide a clear channel importance ranking ^12^. However, it was possible to infer from their work that adding CD4, CD56, and CD8 can improve the classification performance significantly. This is the same combination as suggested by PXPermute. Apart from the current datasets, PXPermute can potentially be used on other data modalities such as multiplex IF images (mIF) ^32–34^ and multiplexed protein maps ^35,36^, where the data includes multiple fluorescent stainings with complex morphologies.

To utilize PXPermute in lab conditions, we recommend performing a small experiment with all the possible stainings and applying PXPermute to the data. After the post-hoc analysis, the main experiment can be done using the selected channels.

A potential limitation in this work is the need to repeat the execution of PXPermute multiple times to enhance its robustness. This iterative process can be time-consuming depending on the dataset size and the number of channels involved. However, it is worth noting that PXPermute has been designed to support parallelization, enabling significant acceleration of the calculations and mitigating this limitation.

In summary, PXPermute is the first method that systematically studies channel importance and can potentially lead to optimizing the workflow of biologists in the lab. PXPermute can be implemented for panel optimization and benefit biologists in their lab work.

## Acknowledgments

We thank Nikolaos Kosmas Chlis for inspiring the project and Katharina Essig for the discussions.

## Data and software availability

We used publicly available data. All the links and required information is provided in the Methods section. The code and instructional notebooks on how to run the code and built analysis pipelines can be found here: https://github.com/marrlab/pxpermute

## Author contributions

SSB & AK implemented code and conducted experiments with support of DW. SSB, AK, DW, and CM wrote the manuscript with FS and SK help. SSB created figures and the main storyline with DW, AK and CM. FS and SK helped with the manuscript narrative and editing. CM supervised the study. All authors have read and approved the manuscript.

## Materials and Methods

### Data sets

All data studied in this work was acquired through imaging flow cytometry. For model training and testing, brightfield, as well as fluorescent channels, were used. In the datasets with a strong imbalance, the training set was oversampled by randomly selecting indices from minority classes with replacement. All images were rescaled to 0 and 1 using the minimum and maximum of the datasets.

### Apoptotic Cells dataset

This dataset was published by Kranich et al. ^18^, who not only solved the binary classification task (apoptotic vs. non-apoptotic cell) but also studied the channel importance directly. Each class is represented by only two channels, one fluorescent (Fig. 2 a). The images were cropped to 32×32. The dataset was accessed here: https://github.com/theislab/dali/tree/master/data

### Synapse formation dataset

Shetab Boushehri and Essig et al. published this dataset ^10,21^ to study the process of synapse formation using IFC. Their dataset includes nearly 2,8 million images containing T cells and B-LCL cells or their conjugates; only 5,221 are labeled by an expert. Each image contains eight channels containing brightfield (BF, stain-free), antibody (Ab, fluorescent), CD18 (fluorescent), F-actin (fluorescent), MHCII (fluorescent), CD3 (fluorescent), P-CD3ζ (fluorescent), and Live/Dead (fluorescent) (Fig. 2b). This annotated subset only contains images of live cells with no antibody. Therefore, Ab and Live/Dead channels are redundant. Moreover, they showed that for the classification of synapses, the most important channels are MHCII, CD3, and P-CD3ζ. Since the images are in different sizes, they have been padded to 128×128 to prepare them for training. The dataset was accessed here: https://doi.org/10.5061/dryad.ht76hdrk7

### White blood cell dataset

The white blood cell dataset ^12^ (WBC) contains 98,013 IFC images, each with twelve channels, which were obtained from two whole blood samples of patients. In each image, a single cell is contained, with eight classes (Fig. 2 c): natural killer (NK) cells (669 samples), neutrophils (18,325 samples), eosinophils (1,537 samples), monocytes (1,282 samples), B cells (889 samples), T cells CD8+ (1,788 samples) and CD4+ (4,476 samples), natural killer T (NKT) cells (1,028 samples), and a separate class comprising unidentifiable cells (1,286 samples). All images were reshaped to 64×64. The channels comprise two brightfield channels, one dark field channel, and nine stained channels. Lippeveld et al. ^12^ have shown that stain-free data suffices to classify monocytes and neutrophils, but to classify others reliably, the dataset must include stained channels.

The dataset was accessed here: https://cloud.irc.ugent.be/public/index.php/s/assnP3Z2FjTbztc

### Classification Models

In this work, we analyzed a commonly used deep learning classifier. The models’ decision-making process cannot be interpreted without applying explainability methods. We selected ResNet18 (16) from all image classification models due to its good performance reported in the previous works ^10,12,24^. We used weights pre-trained on ImageNet ^23^ implemented in the PyTorch package ^37^. To utilize the scikit-learn ^38^ pipeline for the deep learning model, we used the model wrapper from the Skorch library ^39^. The pipeline comprises the data transformation step, which includes normalizing samples with 1st and 99th percentile of images for each dataset, random vertical, horizontal flip, and random noise. To evaluate and compare model accuracy, we have used the F1-score. The model was trained with the cross-entropy loss and AdamW optimizer, with a batch size of 128. The learning rate was set to 0.001 when initializing a new model before starting the training and was kept decreasing by a factor of 0.5 if there was no improvement in the F1-macro score of the validation set for five epochs straight. We have also applied the early stopping technique abrupting the training if the same metric stays constant ± 0.0001 for 50 epochs.

### Model Interpretation

To the best of the authors’ knowledge, no methods can be directly applied to the pre-trained model to evaluate the channel importance. However, some methods evaluate the importance of a single or set of pixels. Thus, our first approach was to aggregate their results per channel. The aggregation method takes the median of the pixel values per channel.

In this study, we have used the following pixel-wise interpretation methods to analyze the channel importance:

### PxPermutes

Analog to pixel-wise interpretation, channel importance can be estimated via conducting sensitive analysis: measuring changes in the model output caused by changes in input. However, our method permutes pixels in the channel compared to occlusion ^26^, which replaces a specific pixel area with an occluding mask. We avoid violating the critical principle of machine learning, which states that training and test sets must be drawn from the same distribution.

Permuting pixels per channel destroys the structural information contained in contours, edges, or areas. It can still be guaranteed that the degradation in the model performance was not due to the artifacts in the distribution.

PXPermute augments each image channel in a dataset by permuting the pixels of each channel *k* times. For a dataset with *N* images, PXPermute generates *N* * *k* modified multichannel images; see Fig. 1 b. The user can define the parameter *k*: a larger *k* leads to more robust results but requires more computational resources. For a very large *k*, the algorithm’s complexity increases and approaches the brute force search approach, in which the model is retrained and re-evaluated for all possible channel combinations. We test PXPermute on a single cell classification task, measuring the drop in F1-score as our metric for performance evaluation. PXPermute design is inspired by the previous works on feature importance as well as interpretability of CNN models ^26,40,41^, which tries to solve the disadvantages of the other interpretability methods and combine all the strengths, such as independence on the model architecture, preserving the same train/test distribution, and simplicity in one method.

### Occlusion

Zeiler et al. ^26^ introduced the idea of estimating the importance of the part of the scene by replacing it with a grey square and observing the classifier’s output. Such a technique was called sensitive analysis or occlusion sensitivity. Later, Zintgraf et al. initiated removing the information from an image completely and calculating the effect. In addition, the authors proposed optimizing the occluding strategy by introducing the marginalization of the occluded pixel. Finally, the method was studied further and formalized by Samek et al. ^42^: *R*_*i*_ = *f*(*x*)⊙*f*(*x* −(1 − *m* _*i*_)), where *R* is a heatmap, *f* is a classifier function, *m* is an indicator function for removing the patch or feature, and ⊙ denotes the element-wise product. As a result, the heatmap highlights the patches (pixels) stronger if their removal affected the classifier results more. Due to its mechanism, the method is referred to as the perturbation-based forward propagation method.

Despite the simplicity of the concept, the method has its disadvantages. One of them is computational complexity. Every time one pixel or patch is occluded, the output must be recomputed. The computational time can rapidly become infeasible if the input is very large. Another problem of the approach is the saturation effect. It occurs when removing only one patch at a time does not affect the output, but removing multiple patches simultaneously does. This leads to misinterpretation and wrong conclusions. In addition, occluding the test image changes its distribution, which can lead to a corrupted interpretation.

### Guided Grad-CAM

another way of interpreting a model is backpropagating an important signal from the output towards the input. If occlusion requires perturbing and passing the input forward multiple times, this approach needs to do a backward pass only once, making it computationally efficient. Nevertheless, gradient-based backpropagation methods have drawbacks, including the saturation effect and being designed for convolutional neural networks only.

Guided Grad-CAM is an element-wise product of the results of two other interpretational approaches: Guided Backpropagation and Grad-CAM ^17^. On the one hand, Grad-CAM uses the last convolutional layer’s gradient information to evaluate each neuron’s importance for a specific class. On the other hand, Guided Backpropagation isn’t class-discriminative but rather highlights details of an image by visualizing gradients in high resolution. Thus, fusing both methods results in high-resolution class-discriminative saliency maps.

### DeepLift

with this method, authors addressed the saturation problem by considering gradient-based interpretation approaches ^27^. Instead of propagating the gradients, this approach suggests propagating the difference from a reference. For example, the reference input can be blurred images or images containing only the background color. Consequently, the method’s output strongly depends on the definition of reference input, which requires data or domain knowledge.

### Layer-wise relevance propagation (LRP)

the idea is to calculate the relevance of input features to the particular prediction ^29^. The relevance is propagated from the output back to the input layers by distributing the output value into relevance scores for each underlying neuron sequentially based on the model’s weights and activations. So that the neuron with a positive contribution receives proportionally a bigger relevance score. The method applies the set of propagation rules depending on the activations and layers, making this method more difficult with the implementation and more computation expensive if applied to a large-scale model.

---

**Supplementary Figure 1:**
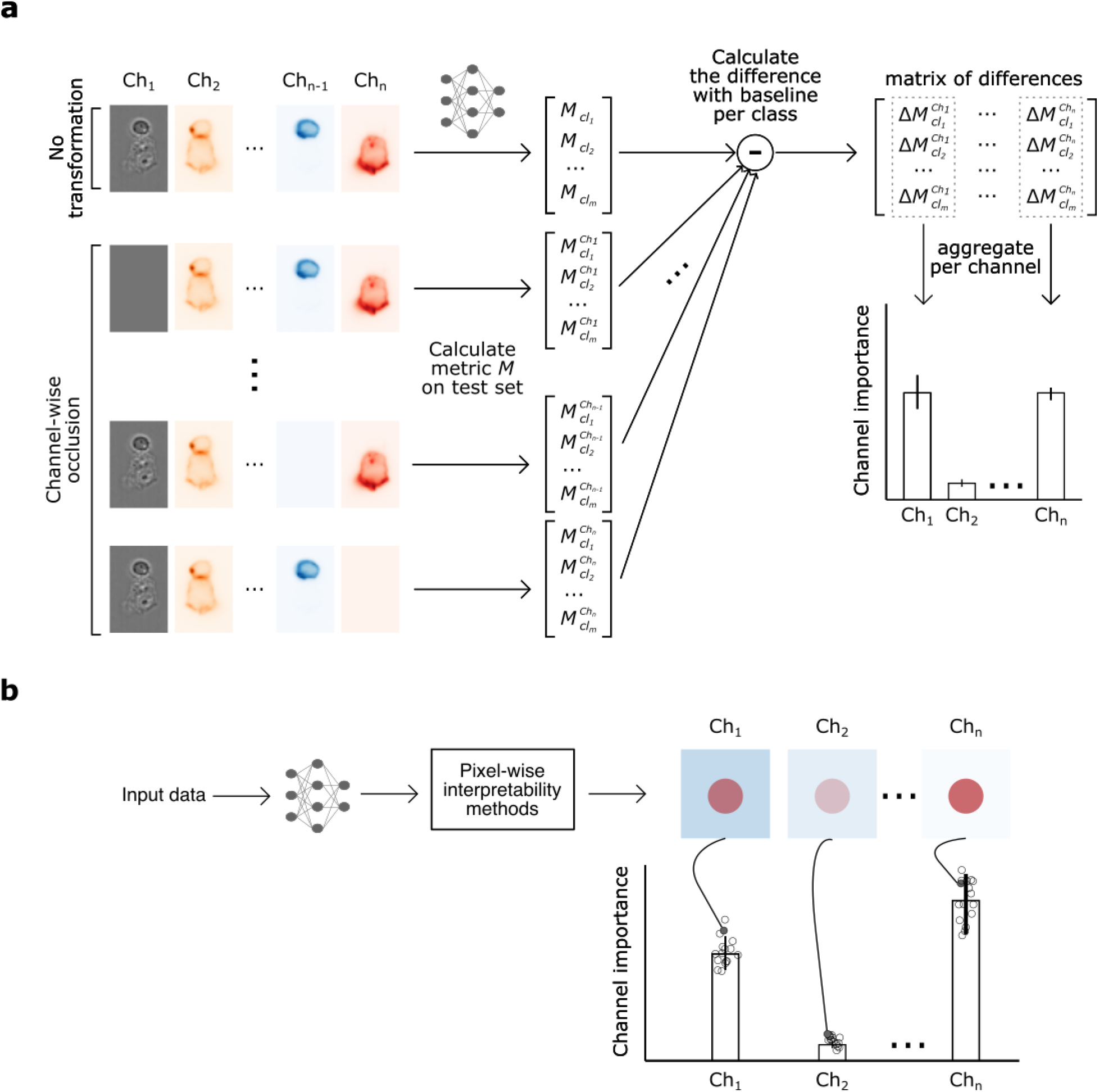
Baseline methods for channel importance measurement and comparison with PXPermute. **a**. Schematic of channel-wise occlusion. In the first step, a performance metric *M*_*cl*_ (such as accuracy or F1-score) is calculated per class *cl*. Then each channel *ch* is occluded with 0, and the performance per class *cl* is calculated as 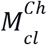 Finally, the difference between the original performance and the permuted one is calculated. These differences are averaged and yield the channel importance **b**. Adaptation of pixel-wise interpretability method for channel importance. For each method, the pixel importance for each sample is calculated. Then the median of the pixel importance per channel is calculated. The aggregation of these values over the whole dataset is considered as channel importance.

**Supplementary Figure 2:**
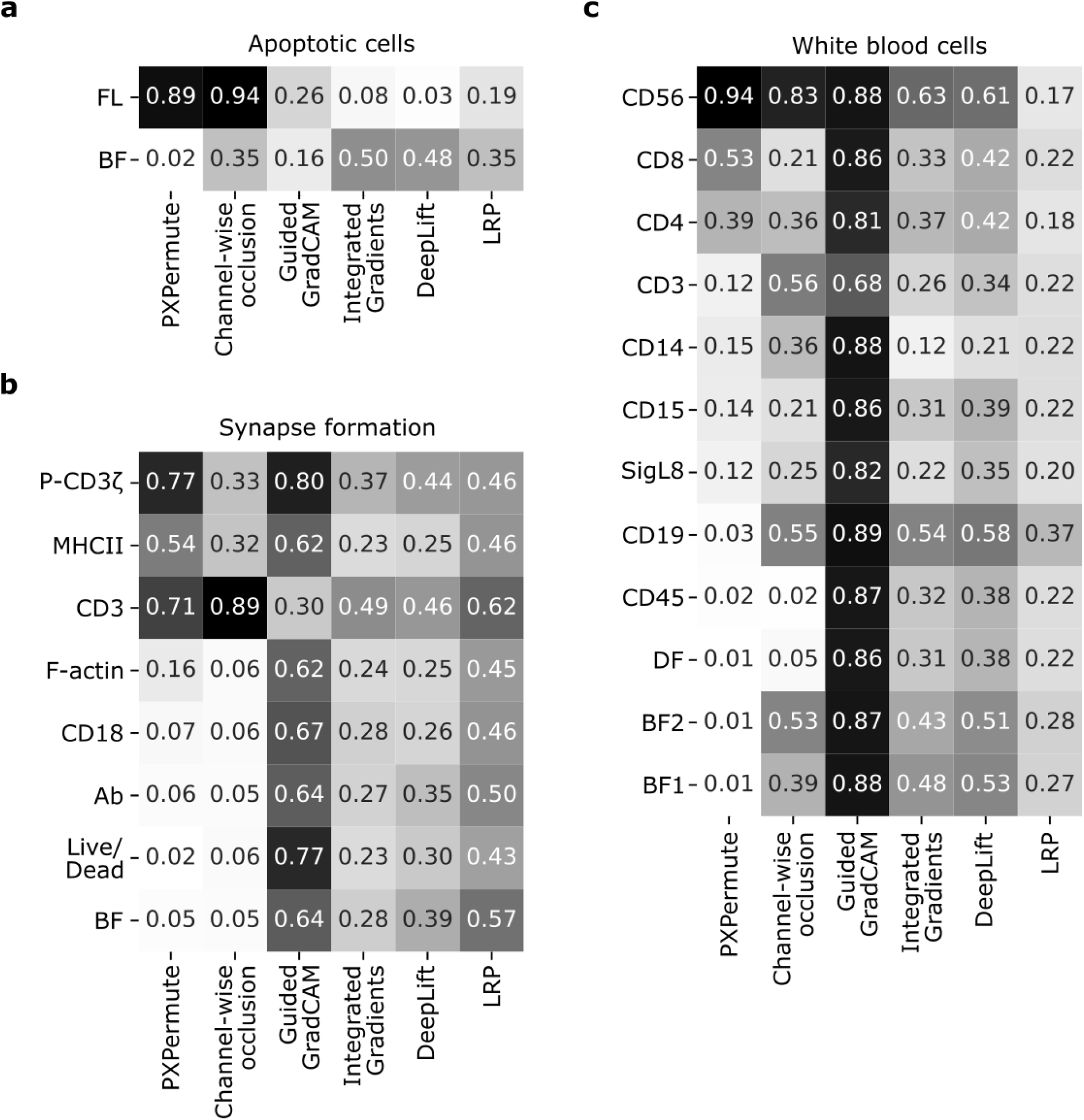
Average channel importance per method. Average value of normalized feature importance for each channel and method per dataset. The values in the heatmaps show the average values in Fig. 3.

## References

1. Rane, A.S., Rutkauskaite, J., deMello, A., and Stavrakis, S. (2017). High-Throughput Multi-parametric Imaging Flow Cytometry. Chem 3, 588–602. 10.1016/j.chempr.2017.08.005.

2. Barteneva, N.S., and Vorobjev, I.A. (2016). Imaging flow cytometry. J. Immunol. Methods.

3. Doan, M., Vorobjev, I., Rees, P., Filby, A., Wolkenhauer, O., Goldfeld, A.E., Lieberman, J., Barteneva, N., Carpenter, A.E., and Hennig, H. (2018). Diagnostic Potential of Imaging Flow Cytometry. Trends in Biotechnology 36, 649–652. 10.1016/j.tibtech.2017.12.008.

4. Barteneva, N.S., Fasler-Kan, E., and Vorobjev, I.A. (2012). Imaging Flow Cytometry: Coping with Heterogeneity in Biological Systems. J. Histochem. Cytochem. 60, 723–733. 10.1369/0022155412453052.

5. Blasi, T., Hennig, H., Summers, H.D., Theis, F.J., Cerveira, J., Patterson, J.O., Davies, D., Filby, A., Carpenter, A.E., and Rees, P. (2016). Label-free cell cycle analysis for high-throughput imaging flow cytometry. Nat. Commun. 7, 10256. 10.1038/ncomms10256.

6. Eulenberg, P., Köhler, N., Blasi, T., Filby, A., Carpenter, A.E., Rees, P., Theis, F.J., and Wolf, F.A. (2017). Reconstructing cell cycle and disease progression using deep learning. Nat. Commun. 8, 463. 10.1038/s41467-017-00623-3.

7. Chlis, N.-K., Rausch, L., Brocker, T., Kranich, J., and Theis, F.J. (2020). Predicting single-cell gene expression profiles of imaging flow cytometry data with machine learning. Nucleic Acids Res. 48, 11335–11346. 10.1093/nar/gkaa926.

8. Lee, K.C.M., Wang, M., Cheah, K.S.E., Chan, G.C.F., So, H.K.H., Wong, K.K.Y., and Tsia, K.K. (2019). Quantitative Phase Imaging Flow Cytometry for Ultra-Large-Scale Single-Cell Biophysical Phenotyping. Cytometry A 95, 510–520. 10.1002/cyto.a.23765.

9. McLaughlin, B.E., Baumgarth, N., Bigos, M., Roederer, M., De Rosa, S.C., Altman, J.D., Nixon, D.F., Ottinger, J., Oxford, C., Evans, T.G., et al. (2008). Nine-color flow cytometry for accurate measurement of T cell subsets and cytokine responses. Part I: Panel design by an empiric approach. Cytometry A 73, 400–410. 10.1002/cyto.a.20555.

10. Boushehri, S.S., Essig, K., Chlis, N.-K., Herter, S., Bacac, M., Theis, F.J., Glasmacher, E., Marr, C., and Schmich, F. (2022). scifAI: Explainable machine learning for profiling the immunological synapse and functional characterization of therapeutic antibodies. bioRxiv, 2022.10.24.513494. 10.1101/2022.10.24.513494.

11. Hennig, H., Rees, P., Blasi, T., Kamentsky, L., Hung, J., Dao, D., Carpenter, A.E., and Filby, A. (2017). An open-source solution for advanced imaging flow cytometry data analysis using machine learning. Methods 112, 201–210. 10.1016/j.ymeth.2016.08.018.

12. Lippeveld, M., Knill, C., Ladlow, E., Fuller, A., Michaelis, L.J., Saeys, Y., Filby, A., and Peralta, D. (2020). Classification of Human White Blood Cells Using Machine Learning for Stain-Free Imaging Flow Cytometry. Cytometry A 97, 308–319. 10.1002/cyto.a.23920.

13. Ota, S., Sato, I., and Horisaki, R. (2020). Implementing machine learning methods for imaging flow cytometry. Microscopy 69, 61–68. 10.1093/jmicro/dfaa005.

14. Lippeveld, M., Peralta, D., Filby, A., and Saeys, Y. (2022). A scalable, reproducible and open-source pipeline for morphologically profiling image cytometry data. bioRxiv, 2022.10.24.512549. 10.1101/2022.10.24.512549.

15. Timonen, V.A., Kerkelä, E., Impola, U., Penna, L., Partanen, J., Kilpivaara, O., Arvas, M., and Pitkänen, E. (2022). DeepIFC: virtual fluorescent labeling of blood cells in imaging flow cytometry data with deep learning. bioRxiv, 2022.08.10.503433. 10.1101/2022.08.10.503433.

16. Strobl, C., Boulesteix, A.-L., Kneib, T., Augustin, T., and Zeileis, A. (2008). Conditional variable importance for random forests. BMC Bioinformatics 9, 307. 10.1186/1471-2105-9-307.

17. Selvaraju, R.R., Cogswell, M., Das, A., Vedantam, R., Parikh, D., and Batra, D. (2017). Grad-CAM: Visual Explanations from Deep Networks via Gradient-Based Localization. In 2017 IEEE International Conference on Computer Vision (ICCV), pp. 618–626. 10.1109/ICCV.2017.74.

18. Kranich, J., Chlis, N.-K., Rausch, L., Latha, A., Schifferer, M., Kurz, T., Foltyn-Arfa Kia, A., Simons, M., Theis, F.J., and Brocker, T. (2020). In vivo identification of apoptotic and extracellular vesicle-bound live cells using image-based deep learning. J Extracell Vesicles 9, 1792683. 10.1080/20013078.2020.1792683.

19. Comeau, J.W.D., Costantino, S., and Wiseman, P.W. (2006). A guide to accurate fluorescence microscopy colocalization measurements. Biophys. J. 91, 4611–4622. 10.1529/biophysj.106.089441.

20. Aaron, J.S., Taylor, A.B., and Chew, T.-L. (2018). Image co-localization - co-occurrence versus correlation. J. Cell Sci. 131. 10.1242/jcs.211847.

21. Essig, K., Boushehri, S.S., Marr, C., Schmich, F., and Glasmacher, E. (2022). An imaging flow cytometry dataset for profiling the immunological synapse of therapeutic antibodies. 10.5061/dryad.ht76hdrk7.

22. He, K., Zhang, X., Ren, S., and Sun, J. (2016). Deep Residual Learning for Image Recognition. In 2016 IEEE Conference on Computer Vision and Pattern Recognition (CVPR), pp. 770–778. 10.1109/CVPR.2016.90.

23. Deng, J., Dong, W., Socher, R., Li, L.-J., Li, K., and Fei-Fei, L. (2009). ImageNet: A large-scale hierarchical image database. In 2009 IEEE Conference on Computer Vision and Pattern Recognition, pp. 248–255. 10.1109/CVPR.2009.5206848.

24. Shetab Boushehri, S., Qasim, A.B., Waibel, D., Schmich, F., and Marr, C. (2022). Systematic Comparison of Incomplete-Supervision Approaches for Biomedical Image Classification. In Artificial Neural Networks and Machine Learning – ICANN 2022 (Springer International Publishing), pp. 355–365. 10.1007/978-3-031-15919-0_30.

25. Xie, Y., and Richmond, D. (2019). Pre-training on Grayscale ImageNet Improves Medical Image Classification. In Computer Vision – ECCV 2018 Workshops (Springer International Publishing), pp. 476–484. 10.1007/978-3-030-11024-6_37.

26. Zeiler, M.D., and Fergus, R. (2014). Visualizing and Understanding Convolutional Networks. In Computer Vision – ECCV 2014 (Springer International Publishing), pp. 818–833. 10.1007/978-3-319-10590-1_53.

27. Shrikumar, A., Greenside, P., and Kundaje, A. (06--11 Aug 2017). Learning Important Features Through Propagating Activation Differences. In Proceedings of the 34th International Conference on Machine Learning Proceedings of Machine Learning Research., D. Precup and Y. W. Teh, eds. (PMLR), pp. 3145–3153.

28. Sundararajan, M., Taly, A., and Yan, Q. (06--11 Aug 2017). Axiomatic Attribution for Deep Networks. In Proceedings of the 34th International Conference on Machine Learning Proceedings of Machine Learning Research., D. Precup and Y. W. Teh, eds. (PMLR), pp. 3319–3328.

29. Bach, S., Binder, A., Montavon, G., Klauschen, F., Müller, K.-R., and Samek, W. (2015). On Pixel-Wise Explanations for Non-Linear Classifier Decisions by Layer-Wise Relevance Propagation. PLoS One 10, e0130140. 10.1371/journal.pone.0130140.

30. Teng, Q., Liu, Z., Song, Y., Han, K., and Lu, Y. (2022). A survey on the interpretability of deep learning in medical diagnosis. Multimed Syst 28, 2335–2355. 10.1007/s00530-022-00960-4.

31. Hooker, S., Erhan, D., Kindermans, P.-J., and Kim, B. (2018). A benchmark for interpretability methods in deep neural networks. arXiv [cs.LG].

32. Eng, J., Bucher, E., Hu, Z., Zheng, T., Gibbs, S.L., Chin, K., and Gray, J.W. (2022). A framework for multiplex imaging optimization and reproducible analysis. Commun Biol 5, 438. 10.1038/s42003-022-03368-y.

33. Lin, J.-R., Izar, B., Wang, S., Yapp, C., Mei, S., Shah, P.M., Santagata, S., and Sorger, P.K. (2018). Highly multiplexed immunofluorescence imaging of human tissues and tumors using t-CyCIF and conventional optical microscopes. Elife 7. 10.7554/eLife.31657.

34. Rojas, F., Hernandez, S., Lazcano, R., Laberiano-Fernandez, C., and Parra, E.R. (2022). Multiplex Immunofluorescence and the Digital Image Analysis Workflow for Evaluation of the Tumor Immune Environment in Translational Research. Front. Oncol. 12, 889886. 10.3389/fonc.2022.889886.

35. Spitzer, H., Berry, S., Donoghoe, M., Pelkmans, L., and Theis, F.J. (2022). Learning consistent subcellular landmarks to quantify changes in multiplexed protein maps. bioRxiv, 2022.05.07.490900. 10.1101/2022.05.07.490900.

36. Gut, G., Herrmann, M.D., and Pelkmans, L. (2018). Multiplexed protein maps link subcellular organization to cellular states. Science 361. 10.1126/science.aar7042.

37. Paszke, A., Gross, S., Massa, F., Lerer, A., Bradbury, J., Chanan, G., Killeen, T., Lin, Z., Gimelshein, N., Antiga, L., et al. (2019). PyTorch: An imperative style, high-performance deep learning library. arXiv [cs.LG].

38. Pedregosa, F., Varoquaux, G., Gramfort, A., Michel, V., Thirion, B., Grisel, O., Blondel, M., Müller, A., Nothman, J., Louppe, G., et al. (2012). Scikit-learn: Machine Learning in Python. arXiv [cs.LG], 2825–2830.

39. Skorch documentation — skorch 0.12.1 documentation https://skorch.readthedocs.io/en/stable/.

40. Altmann, A., Toloşi, L., Sander, O., and Lengauer, T. (2010). Permutation importance: a corrected feature importance measure. Bioinformatics 26, 1340–1347. 10.1093/bioinformatics/btq134.

41. Breiman, L. (2001). Random Forests. Mach. Learn. 45, 5–32. 10.1023/A:1010933404324.

42. Samek, W., Binder, A., Montavon, G., Lapuschkin, S., and Muller, K.-R. (2017). Evaluating the Visualization of What a Deep Neural Network Has Learned. IEEE Trans Neural Netw Learn Syst 28, 2660–2673. 10.1109/TNNLS.2016.2599820.

